# Local Accumulation of Axonal Mitochondria in the Optic Nerve Glial Lamina Precedes Myelination

**DOI:** 10.1101/2021.02.25.432924

**Authors:** Samantha J. Wilkison, Cora L. Bright, Ricardo Vancini, Daniel J. Song, Howard M. Bomze, Romain Cartoni

## Abstract

Mitochondria are essential for neurons and must be optimally distributed along their axon to fulfil local functions. A high density of mitochondria has been observed in retinal ganglion cell (RGC) axons of an unmyelinated region of the optic nerve, called the glial lamina (GL) in mouse (lamina cribrosa in human). In glaucoma, the world’s leading cause of irreversible blindness, the GL is the epicenter of RGC degeneration and is connected to mitochondrial dysfunction. It is generally accepted that the local accumulation of mitochondria in the GL is established due to the higher energy requirement of unmyelinated axons. Here we revisit the connection between mitochondrial positioning and myelin in RGC axons. We show that the high density of mitochondria in the GL is restricted to larger axons and is established before myelination. Thus, contrary to a longstanding belief in the field, the myelination pattern is not responsible for the establishment of the local accumulation of mitochondria in GL axons. Our findings open new research avenues likely critical to understanding the pathophysiology of glaucoma.

## Introduction

How neurons allocate resources along the axon is critical for their physiology and survival. Chief among these cellular resources are mitochondria, which must be distributed to specific axonal regions to accomplish local functions. Neurons rely heavily on mitochondria for ATP production (Ames, 2000; Zala et al., 2013), calcium buffering (Rizzuto et al., 2012) and reactive oxygen species management (Angelova and Abramov, 2018). Due to the size of their axonal projection, these important functions must be fulfilled locally in critical area along the axon (MacAskill and Kittler, 2010; Misgeld and Schwarz, 2017). For example, during development, cortical neurons capture mitochondria at future axonal branch point (Courchet et al., 2013). When axons are growing or re-growing after an injury, mitochondria are re-localized to provide the growth cone with the energy needed (Misgeld et al., 2007; Morris and Hollenbeck, 1993; Zhou et al., 2016). Therefore, it is important to have a clear picture of how mitochondria are distributed in specific axonal region and to understand what factors regulates this positioning. RGCs are neurons from the central nervous system located in the inner retina that project their axons in the optic nerve to reach their targets in the brain via the optic track. RGC axons are heterogeneously myelinated: In the proximal region of the optic nerve head called the glial lamina (GL) in mouse and lamina cribrosa in human they are unmyelinated whereas the distal retrolaminar region (RL) of the nerve is fully myelinated (Fig.1A). The physiological function of this local lack of myelination is unclear but the importance of this optic nerve region is evidenced by the well-established vulnerability of the GL in glaucoma, the leading cause of irreversible blindness worldwide, in which an early insult to axons in the GL triggers RGC degeneration (Nickells et al., 2012). Of significance, mitochondrial dysfunction has been shown to be important to the pathophysiological mechanism of glaucoma (Chrysostomou et al., 2013; Osborne et al., 2016; Williams et al., 2017). Seminal studies have described a higher abundance of axonal mitochondria in the GL compared to the RL (Barron et al., 2004; Bristow et al., 2002; Yu Wai Man et al., 2005; Yu et al., 2013). Despite the relevance of this axonal region in the context of glaucoma little is known about the mitochondrial landscape in axons of the GL both in adult and during development. Since unmyelinated axons requires more ATP to propagate axon potential (Perge et al., 2009), it has long been accepted that the axonal mitochondrial accumulation in the GL is established in response to the higher energy requirement of unmyelinated axons (Barron et al., 2004; Bristow et al., 2002; Yu Wai Man et al., 2005; Yu et al., 2013). However, direct evidence for this causal relationship is lacking. Here, we sought to investigate this relationship by imaging the positioning of mitochondria in mouse RGC axons throughout different myelination stages. Using two imaging techniques namely serial block face scanning electron microscopy (SBF-SEM) and light sheet fluorescent microscopy (LSFM), we found that the high abundance of mitochondria in GL axons is observed specifically in larger axons independently of the myelination status. Furthermore, we demonstrated that this mitochondrial accumulation was established early during development between P5 and P6-P8 a stage preceding RGC axons myelination.

## Materials and Methods

### Animals

Experiments using mice were approved by the Duke University Institutional Animal Care and Use Committee (protocols A194-20-10). The mice were housed under a 12 h light-dark cycle with ad lib access to food and water. Heat and humidity were maintained within the parameters specified in the National Institute of Health Guide for the Care and Use of Laboratory Animals. Experimental procedures were also consistent with this Guide. STOP^f/f^-mitoEGFP;Vglut2-Cre (mitoRGC) mice used for LSFM study were generating by crossing STOP^f/f^-mitoEGFP with Vglut2-Cre mice both purchased from The Jackson Laboratory (stock 021429 and 028863 respectively). Wild-type C57BL/6J mice (The Jackson Laboratory stock 000664) were used for the SBF-SEM study.

### LSFM Samples Preparation

The optic nerves underwent immunohistochemistry staining protocol adapted from McKey et al. (2020) (McKey et al., 2020). Adult mice were transcardially perfused using with 1x PBS and 4% paraformaldehyde (PFA) and neonatal pups heads were drop fixed for 2 hours in 4% PFA. Optic nerves were then dissected from the surface of the optic nerve head to the optic chiasm and underwent a sequential dehydration sequence into 100% methanol. Samples were left in a 66% Dichloromethane, 33% methanol mixture overnight. After washing with 100% methanol, the samples were bleached with 5% hydrogen peroxide and 95% methanol then sequentially rehydrated and permeabilized in PTx.2, glycine and DMSO. Nerves were then transferred into blocking solution overnight (PTx.2, 1.5% donkey serum, 10% DMSO) and put into primary antibody (PTwH, 5% DMSO, 3% donkey serum, Goat anti-GFP, Abcam ab5450, 1:1000) for 10 days at 37 degrees Celsius. Following three washes of PTwH, samples were incubated with secondary antibody (PTwH, 3% donkey serum, Cy3-conjugated Donkey Anti-Goat, Jackson ImmunoResearch Labs 705-165-147, 1:1000) overnight. Samples were then washed three times, with PTwH. Adult optic nerves were dehydrated sequentially (25%, 50%, 75%, 100% MeOH/PBS respectively). Once dehydrated, samples are incubated with 66% DCM and 33% methanol, washed with 100% DCM and left in 100% Dibenzyl ether overnight. Samples were again rehydrated sequentially (100%, 75%, 50%, 25% MeOH/PBS respectively) and put into Cubic 1 solution overnight (25% urea, 25% of 80% quadrol, 15% Triton X-100, diH2O). Samples were then incubated in Cubic 2 solution (25% urea, 50% sucrose, 10% triethanolamine, diH2O) over night. Adult and neonatal optic nerves were embedded in 1.8% agarose using a 1 mL syringe. Once agarose has solidified, samples were expelled out of the syringe so they can hang in Cubic 2 solution overnight and imaged the next day.

### LSFM Acquisition

Image collection was performed on a Zeiss Lightsheet Z.1equipped with CLARITY 20x objective, Nd =1.45 and NA=1.0, and pco.Edge sCMOS cameras (dual). Illumination was dual sided with 10x NA 0.2 objectives and 561 nm diode pump solid state laser (set at 5 to 6% of power) via LBF 405/488/561/638 quad dichroic and a Zeiss ET600/50 nm emission filters. Cleared optic nerves were mounted from a syringe into an imaging sample holder and submerged in Cubic 2 solution prepared fresh with a average refractive index of 1.45. Depending on the data set, the image pixel size ranged from 0.28-0.38 μm and Z-stacks were collected with a Z-step ranging from 0.585-1.24 μm. The system was controlled by Zeiss Zen 9.2.0.0 2014 SP1 (black edition) for lightsheet.

### LSFM Image Analysis

The sequence of z-stack images was opened in Fiji. The background of each image was then measured using the rectangle tool and the minimum value of the pixel intensity histogram was subtracted from each slice. From the projected z-stack image, the pixel intensity along the nerve was measured by drawing a line (line tool) covering the entire nerve.

### Serial Block Face Scanning Electron Microscopy

Wild Type (C57BL/6J) mice were perfused using Ringer Solution for 2 minutes (sodium chloride 123 mM), calcium chloride 1.5 mM, potassium chloride 4.96 mM pH 7.3-7.4) followed by a fixative solution (0.15 M cacodylate, 2.5% glutaraldehyde, 2% paraformaldehyde and 2 mM calcium chloride) for 2 minutes. After fixation, the samples underwent a heavy metal staining protocol adapted from Deerinck et al. (14). Briefly, samples were washed in 0.1 M sodium cacodylate pH 7.4 and then transferred for 1h at room temperature into a solution of 1.6 % potassium ferrocyanide containing 2% osmium tetroxide buffered with 0.1 M sodium cacodylate followed by 30min in fresh and filtered 10% thiocarbohydrazide (TCH) solution. Samples were then washed in distilled water and incubated a secondary 2% aqueous osmium tetroxide incubation for 30min. The samples were then placed in 1% aqueous uranyl acetate at 4°C overnight, washed in distilled water, and placed in freshly prepared lead aspartate solution (0.066g of lead nitrate in 10ml 0.03M of aspartic acid solution) for 30min at 60°C. The samples were dehydrated in a graded series of with cold ethanol, from 25% to 100% and then infiltrated with increasing concentrations of Durcupan resin in ethanol with several exchanges of 100% resin. The samples were finally embedded in 100% resin and allowed to polymerize at 60ଌ for 48 hours. The tissue samples embedded in resin were manually trimmed with a razor blade to expose the tissue on their surfaces and then glued onto an aluminum SBF-SEM rivet with conductive epoxy (SPI Conductive Silver Epoxy) with the exposed tissue down. Specimens on the rivet were further trimmed by hand with a razor blade to as small a size as possible (about 0.5 mm), and block face was trimmed with a glass knife. Once tissue was exposed, semi thin sections 0.5 μm were cut and placed on a glass slide; they were stained with toluidine blue and viewed under a light microscope to check tissue orientation, condition, and correct localization. The rivet with the sample was then sputter coated with gold-palladium to ensure electrical conductivity of the tissue edges with the stub.

### Acquisition of SBF-SEM image stacks

The image stacks were acquired in an automated fashion by using a high-resolution field emission scanning electron microscope (SEM) (Merlin - Carl Zeiss, Germany) equipped with a 3View system (Gatan Inc., Pleasanton, CA, USA), and a back-scattered electron detector. Digital Micrograph software (Gatan Inc.) was used to adjust the imaging conditions and slicing parameters. The SEM was operated in the high-resolution mode with an acceleration voltage of 2 kV current mode and in the high-vacuum mode. All images were taken at settings of 80 pA, 2s dwell time, and 5-7 nm pixel size. Between 50 and 60 sections were obtained at 60nm thickness through 3μm deep, covering fields of view of 75μm x 75 μm (neonatal optic nerves) and 100 μm x 100 μm (adult optic nerve).

### SBF-SEM image analysis

Raw dm4 files were converted to 8bit tiff images and analyzed using Fiji. To assess the axonal size-dependent mitochondrial content, multiple regions of interest were randomly chosen from the original image. The perimeter of each axon analyzed was manually determined using the tracing tool, and the number of mitochondria was counted using the counting tool. Each axon and mitochondrion were given a unique identification number for each image which allowed tracing back to any measurement. For the G ratio, measurement of myelinated axons, the inner axonal area was divided by the outer total axonal area. For unmyelinated axons, the outer total axonal area was divided by itself.

### Statistical analysis

Statistical analysis was performed using GraphPad Prism software. Specific sample size, statistical test and p values for each experiment are given in the appropriate figure legends. p value less than 0.05 was considered significant.

## Results

We first used SBF-SEM to image optic nerve cross-sections obtained from the GL and from the RL, respectively within 150 μm and over 300 μm from the surface of the optic nerve head (Fig. 1A). We quantified the mitochondrial content of axons in the GL and RL in which at least one mitochondrion was identified since the percentage of axons with mitochondria was not higher in the GL comparing to RL in any age we studied (data not shown). In accordance with published studies (Barron et al., 2004; Bristow et al., 2002; Yu Wai Man et al., 2005; Yu et al., 2013), we observed an increase in the mitochondrial content of axons of the GL region compared to RL region (Fig.1B-C). This significant but modest difference led us to hypothesize that the proximal mitochondrial accumulation might not occur uniformly across the axonal population. Although the size distribution of axons differs between the GL and the RL, likely due to the astrocytic web that bundles the axons of the GL, both regions have a wide range of axonal size (from 0.1 μm^2^ to over 2 μm^2^, data not shown) which likely influences mitochondrial content. We therefore analyzed the number of mitochondria per axon across the range of axonal area and demonstrated that the high mitochondrial content in the GL was restricted to larger axons (Fig. 1D-E). Axons with an area above 1 μm^2^ show a 51.5% increase in mitochondrial abundance in the GL (GL: 2.47 +/- 0.09 vs RL: 1.63 +/- 0.03 respectively) whereas axons smaller than 1 μm^2^ show similar mitochondrial density in the GL and RL (GL: 1.18 +/- 0.01 vs RL: 1.11 +/- 0.01) (Fig.1E).

**Figure 1.**
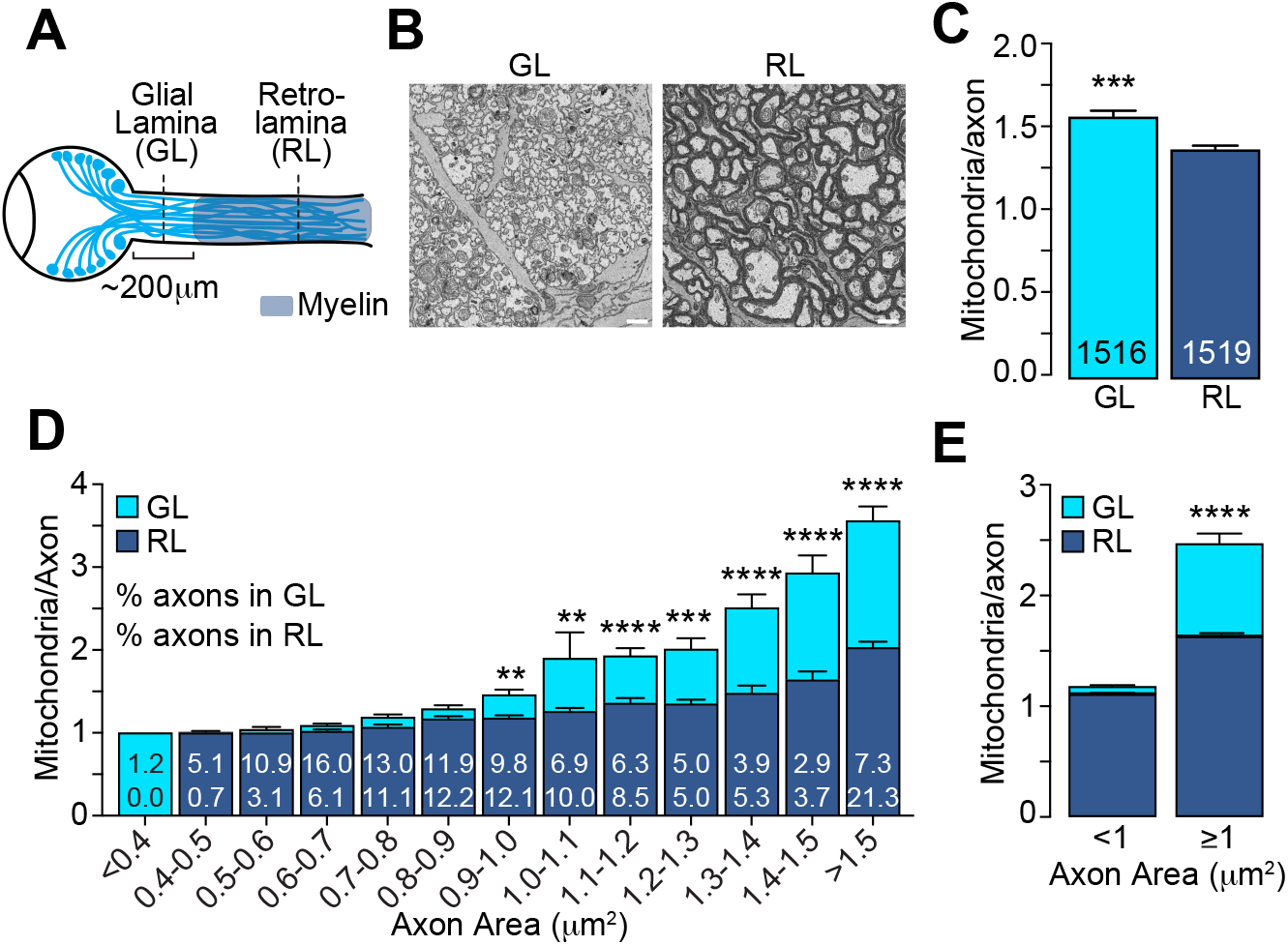
The mitochondrial accumulation at the glial lamina is specific to larger axons. (A) Schematic of the adult mouse optic nerve. (B) Representative electron micrographs of optic nerve cross-sections obtained at the myelinated GL and unmyelinated RL regions. Scale bar=1 μm. (C) Average number of mitochondria per axon in the GL and RL regions. *** p=0.0003, Mann Whitney test. The total number of axons are indicated in the bars; n=3 mice. (D) GL and RL average number of mitochondria per axon as a function of axonal area. The numbers within each superimposed bar represent the percentage of total axons falling within each area range for the GL (top) and RL (bottom). **p<0.002, ***p<0.0006, ****p<0.0001, Kruskal-Wallis test and Dunn’s multiple comparisons. (E) Average number of mitochondria per axon in the GL and RL for axons with areas smaller or bigger than 1 μm^2^. ****p<0.0001, Kruskal-Wallis test and Dunn’s multiple comparisons. Error bars=SEM.

We next tested the paradigm that the absence of myelin around axons in the GL directly induces the local increase in their mitochondrial content. This model predicts that the difference in mitochondrial content between axons in the GL and RL regions should only be apparent after myelination. Therefore, we assessed the mitochondrial density in axons in GL and RL regions of neonatal mice, which have not yet undergone myelination. To efficiently screen multiple neonatal stages, we used LSFM to image the mitochondrial positioning in RGC axons in the intact optic nerve of a Vglut2-Cre; STOP^f/f^-mitoEGFP (mitoRGC) transgenic mouse that expresses a mitochondria-targeted EGFP in early embryonic post-mitotic RGCs. As expected, and consistent with our results shown in Figure 1, we observed a mitochondrial enrichment in the first 200 μm of the adult mitoRGC optic nerve, corresponding to the GL (Fig. 2A left, 2B and Movie S1). Optical cross sections comparing GL and RL confirmed a non-homogenous axonal distribution of mitochondria between these two regions (Fig. 2A, lower panels). MitoRGC optic nerves incubated only with the secondary antibody did not show any signal over background (Fig. 2A right, 2D right). Surprisingly, in unmyelinated optic nerves of neonatal mitoRGC mice (P6 to P9) (Fig. 2C) we also observed a strong mitochondrial enrichment in proximal axons (Fig. 2D, 2E and Movie S2). However, this proximal mitochondrial accumulation was absent in P3 to P5 nerves (Fig. 2F, 2G and Movie S3) in which the level of mitochondrial staining was similar to the RL of adult and P8 (Fig. 2A and D).

**Figure 2.**
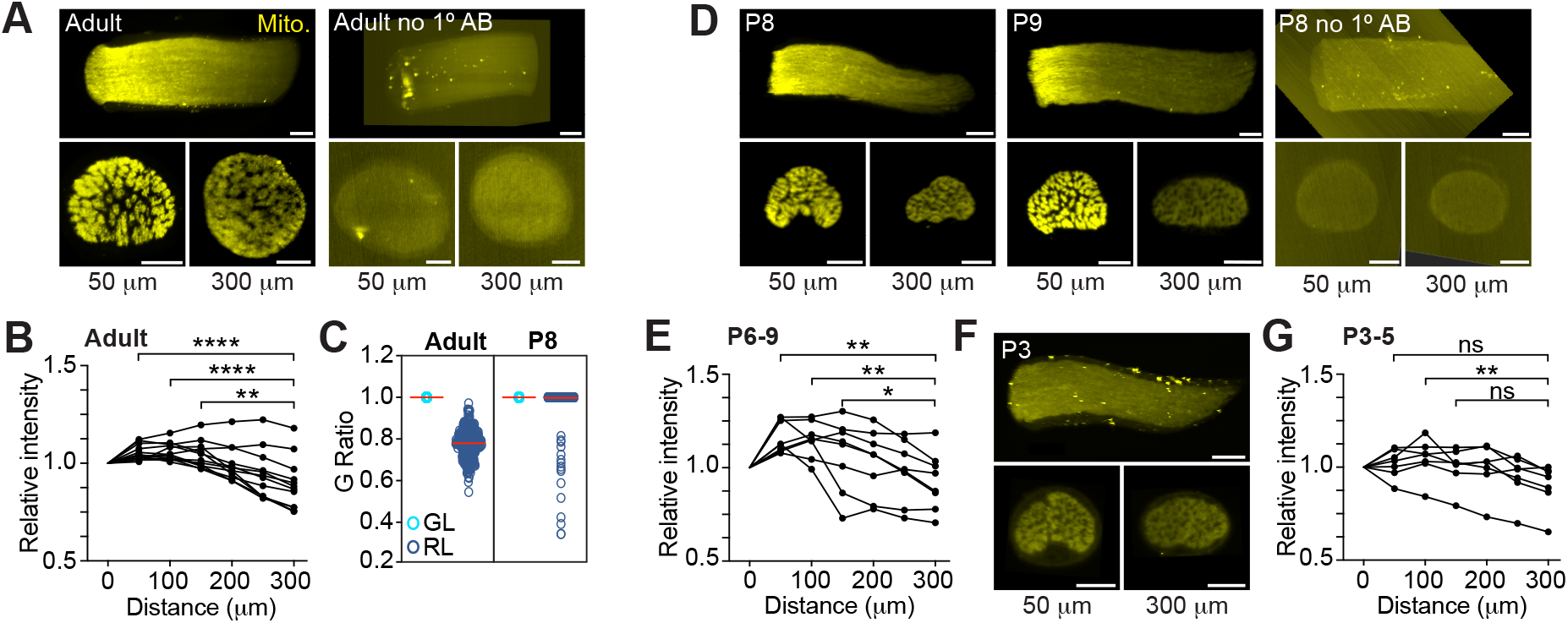
The mitochondrial accumulation in axons of the glial lamina precedes myelination. (A) Representative 3D reconstruction of LSFM images (longitudinal view) of an adult mitoRGC optic nerve (left) and negative control (right). Optical 1.5 μm thick cross-sectional images taken at 50 (GL) and 300 μm (RL) from the surface of the optic nerve head are shown underneath the nerve images. (B) Quantification of the relative fluorescence along the adult mitoRGC optic nerves. **p=0.002, ****p<0.0001, Friedman test and Dunn’s multiple comparisons, n=12 optic nerve, 12 mice. (C) G Ratio of axons in the GL and RL regions in adult and P8 mice. A G ratio less than 1 signifies a myelinated axon. Adult proximal: n=3042 axons, 3 mice, 3 nerves. Adult distal: n=321 axons, 3 mice, 3 nerves. P8 proximal: n=1882 axons, 3 mice, 3 nerves. P8 distal: n=2476 axons, 4 mice, 4 nerves. (D) Representative LSFM images of a P8 and P9 mitoRGC optic nerve and P8 negative control. (E) Quantification of the relative fluorescence along P6 to P9 mitoRGC optic nerves. **p<0.007, *p=0.037, Friedman test and Dunn’s multiple comparisons, n=8 optic nerve, 8 mice. (F) Representative LSFM images of a P3 mitoRGC optic nerve. (G) Quantification of the relative fluorescence along P3 to P5 mitoRGC optic nerves. **p=0.002, ns=non-significant, Friedman test and Dunn’s multiple comparisons, n=7 optic nerve, 7 mice. Scale bar=100 μm.

We next validated this result at single axon resolution using SBF-SEM. We showed that even though 99.7% of the RL axons are unmyelinated at this age (Fig. 2C) we detected an axonal area-dependent increase in mitochondria per axon in the GL of P8 animals (Fig. 3A-D), reminiscent of the adult phenotype (Fig. 1C-E). Axons larger than 0.5 μm^2^ show a 28.7% increase in mitochondrial abundance in the GL (GL: 1.64 +/- 0.11 vs RL: 1.27 +/- 0.06) whereas it remains constant in axons smaller than 0.5 μm^2^ (GL: 1.07 +/- 0.008 vs RL: 1.058 +/- 0.007) (Fig. 3 C and D). Consistent with our results using LSFM, the mitochondrial content was similar in axons of the GL and RL in P5 mice (Fig. 3E-G) suggesting that the local accumulation of mitochondria in large GL axons is developmentally regulated and is set between P5 and P6-P8, a stage that precedes axon myelination in the optic nerve. Overall, our results demonstrate that from P6-P8 large axons of the optic nerve increase their mitochondria abundance in the GL region comparing to the RL whereas small axons keep a constant number of mitochondria per axons in both region (Fig. 4).

**Figure 3.**
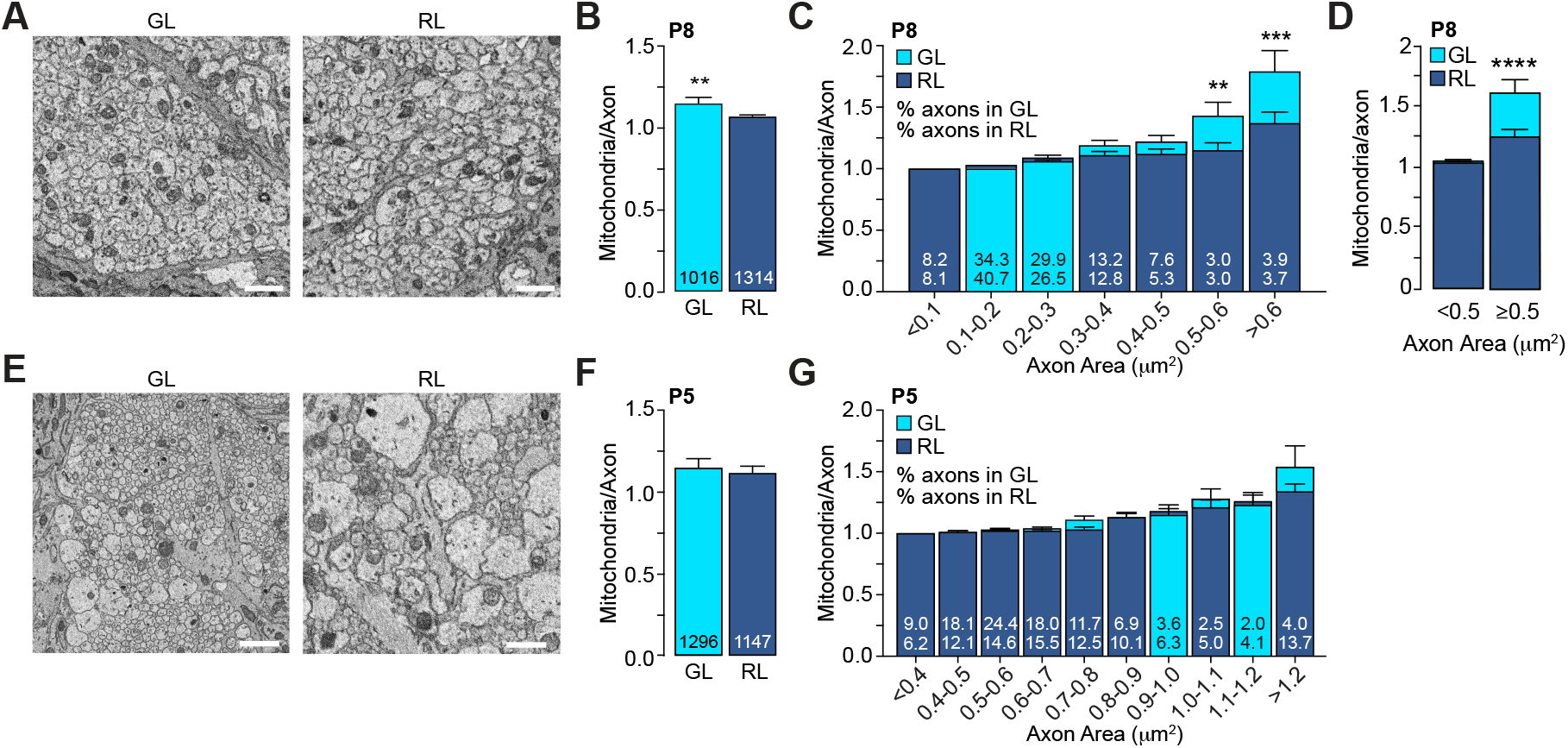
The mitochondrial accumulation in large axons of the glial lamina is established between P5 and P8. (A) Representative electron micrographs of the axons in GL and RL of P8 mice. (B) Average number of mitochondria per axon in axons in GL and RL of P8 mice. The total number of axons is indicated in the bars. GL: n=3 mice, 3 nerves. RL: n= 4 mice, 4 nerves. ***p=0.004, Mann Whitney test. (C) GL and RL average number of mitochondria per axons in P8 mice split by axonal area. The numbers within each superimposed bar represent the percentage of total axons falling within each area range for the GL (top) and RL (bottom). **p=0.006, ***p=0.0002, Kruskal-Wallis test and Dunn’s multiple comparisons. (D) Average number of mitochondria per axon in the GL and RL for axons with areas smaller or bigger than 0.5 μm^2^, ****p<0.0001, Kruskal-Wallis test and Dunn’s multiple comparisons. (E) Representative electron micrographs of the GL and RL axons of P5 mice. (F) Average number of mitochondria per axons in GL and RL of P5 mice. The total number of axons are indicated in the bars. GL: n=4 mice, 4 nerves. RL: n=4 mice, 4 nerves. (G) GL and RL average number of mitochondria per axon in P5 mice split by axonal area. The numbers within each superimposed bar represent the percentage of total axons falling within each area range for the GL (top) and RL (bottom). Scale bar=1μm. Error bars=SEM.

**Figure 4.**
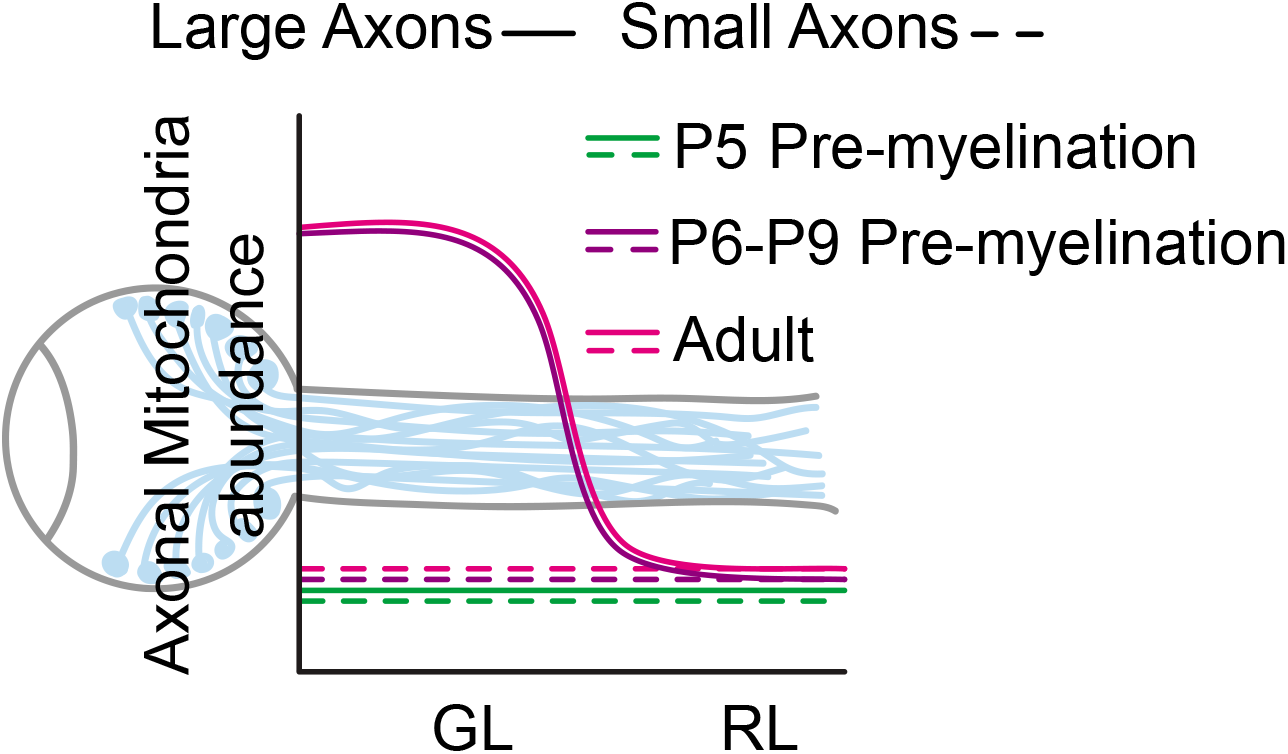
Model of the mitochondrial distribution in RGC axons of the optic nerve GL and RL area.

## Discussion

Using two independent approaches, we revisited an old dogma in neuro-ophthalmology regarding the cause of asymmetrical mitochondria distribution in RGC axons. We provided evidence that the preferential accumulation of mitochondria in axons of the GL is restricted to larger axons and is established between P5 and P6-P8. Since this stage precedes RGC axonal myelination of the optic nerve (Mayoral et al., 2018), this result indicates that the establishment of the mitochondrial enrichment in axons in the GL is not a consequence of the local lack of myelination and occurs in absence of the heterogenous myelination pattern of the optic nerve. We cannot rule out that in the adult optic nerve the persistence of high mitochondrial density in large axons in the GL is functionally related to the lack of myelin, since unmyelinated axons require more energy than their myelinated counterpart (Perge et al., 2009). However, our data clearly indicate that a separate mechanism is required for the initial establishment of the asymmetric mitochondrial distribution along the optic nerve. More work is needed to determine any true functional relationship between axonal mitochondrial distribution and myelination pattern. Remarkably, the axonal size specificity of this local accumulation of mitochondria is present in both neonatal and adult optic nerves; within their own axonal range, only large axons in the GL of adult and P8 show a mitochondria abundance higher than in the RL (Fig. 1 and 3). However, these axons with a mitochondrial accumulation in the GL represent ∼7% of the total axonal population in P8 but ∼50% in adult (Fig. 1D and 3C) suggesting that, as the optic nerve grows, the mitochondrial enrichment in larger axons spread across a broader population of axon. It is well established that unmyelinated axons require more energy than myelinated axons of the same diameter (Perge et al., 2009; 2012). However, our data suggest that smaller axons do not preferentially accumulate mitochondria in unmyelinated regions. This result is consistent with data from adult guinea pig showing that, within the distal optic nerve, the mitochondrial concentration is higher in unmyelinated axons but only in larger axons (Perge et al., 2009). Considerable efforts have been made recently to characterize molecularly the different RGC subtypes (Krieger et al., 2017; Rheaume et al., 2018; Tran et al., 2019) and the function and type specificity of dendritic morphology (Liu and Sanes, 2017; Ran et al., 2020). However, much less is known on the axonal morphology of RGC subtypes, and it is still elusive whether axons of specific size in the optic nerve can be matched to specific RGC subtypes which might reveal common functionality. Therefore, more studies will be needed to decipher if axons accumulating mitochondria proximally in the GL correspond to a combination of RGC subtypes or whether axon size is the only common denominator. Strikingly, the establishment of the local mitochondrial accumulation in large axons of the GL (P6-P8) corresponds to the end of RGC pruning phase, suggesting that the consolidation phase of RGCs and their axons might be coupled with the fine tuning of their mitochondrial landscape (Young, 1984). In glaucoma, RGC death is triggered by an early degeneration of axons in the GL which suggests a local vulnerability likely linked to mitochondria dysfunction (Howell et al., 2007; Williams et al., 2017). This study brings a new perspective on the regulation of the mitochondrial content in axons of this critical region which is highly relevant to the pathophysiology of glaucoma. It also provides new research avenue on the axonal mechanism of mitochondrial positioning.

## Acknowledgments

We thank the Duke University School of Medicine for institutional support and the Whitehead Family Fund (RC), the Research to Prevent Blindness Unrestricted Grant (Duke University) and the Duke University School of Medicine Core Facility Voucher Program (RC). We are grateful to Dr. Carlson and the Duke Light Microscopy Core Facility (National Institute of Health 1S10OD020010-01A1), to Dr. Miller and the Center for Electron Microscopy and Nanoscale Tomography, Department of Pathology, Duke University School of Medicine (National Institute of Health 1S10OD018439). We thank Dr. McKey for critical help with sample preparation for the LSFM, Drs. Sidney M. Gospe III, James V. Alvarez, Nahid Iglesias, Jeremy N. Kay and members of the Cartoni lab for comments on the manuscripts.

## Author Contributions

S.J.W. designed and performed the experiments and analyzed the data. C.L.B. designed and performed experiments and provided assistance in analyzing SBF-SEM data. R.V. provided technical assistance for tissue preparation and performed the SBF-SEM image acquisition. D.J.S, provided assistance on SBF-SEM data analysis and processing. H.M.B. provided technical assistance and intellectual inputs. R.C. designed the experiments, analyzed the data and wrote the manuscript.

## Movies

**Movie S1**. LSFM 3D reconstruction of an adult STOP^f/f^-mitoEGFP; Vglut2-Cre (mitoRGC) optic nerve. Approximatively 0.5 mm of the nerve is shown. In the first frame, the optic nerve head is on the left part of the nerve. The tissue was labelled with an antibody against GFP.

**Movie S2**. LSFM 3D reconstruction of a P8 STOP^f/f^-mitoEGFP; Vglut2-Cre (mitoRGC) optic nerve. Approximatively 0.5 mm of the nerve is shown. In the first frame, the optic nerve head is on the left part of the nerve. The tissue was labelled with an antibody against GFP.

**Movie S3**. LSFM 3D reconstruction of a P5 STOP^f/f^-mitoEGFP; Vglut2-Cre (mitoRGC) optic nerve. Approximatively 0.5 mm of the nerve is shown. In the first frame, the optic nerve head is on the left part of the nerve. The tissue was labelled with an antibody against GFP.

## Notes

### Competing Interest Statement

The authors have declared no competing interest.

